# Analysis of Sexual vs Asexual Reproduction Using Four Measurements

**DOI:** 10.1101/2022.02.05.479258

**Authors:** Feng Lin, Robert J. Lin

**Affiliations:** Department of ECE, Wayne State University, Detroit, MI 48202, USA; Lin Group, Houston, TX 77098, USA

## Abstract

The prevalence of sexual reproduction has long been an outstanding problem of evolutionary biology. In accordance with the mathematical approach employed by past researchers, we propose a mathematical framework to address this problem. We define and derive four measurements, diversity measure DVM, diffusion measure DFM, optimality measure OPM, and survivability measure SVM to compare sexual reproduction with asexual reproduction. We show that DVM increases exponentially in sexual reproduction, while only linearly in asexual reproduction. Hence, sexual reproduction allows species more opportunity to adapt. We also show that DFM is bounded in sexual reproduction and OPM is inversely related to DFM. Thus, sexual reproduction leads to smaller DFM and hence a larger OPM. We further show that SVM is a monotonic increasing function of OPM. Hence, sexual reproduction is better by virtue of producing a more homogeneous population.

## Introduction

Why most species reproduce sexually is a question investigated extensively, but not yet answered satisfactorily in biology (*1–4*). The prevalence of sexual reproduction suggests that there are major benefits provided by this mode of reproduction. The benefits, however, are not well-understood. In many ways, asexual reproduction seems to be a better evolutionary strategy: only one parent is required, and all of the parent’s genes are passed on to its progeny (*5–12*). In a sexual population, the males are unable to produce progeny of their own and females only transfer half their genes to progeny, hence the so called problem of the “two-fold cost of males.” Furthermore, sexual reproduction must also overcome obstacles that do not exist in asexual reproduction. Sexually reproducing organisms must spend a great deal of time and energy to find and attract mates. Also, copulation in sexually reproducing organisms leaves both organisms vulnerable to predation.

Despite these and other major drawbacks to sexual reproduction, it remains a very prevalent form of reproduction in most higher order organisms. Biologists have put forth numerous hypotheses for why sexual reproduction is so prevalent. The main hypotheses to explain sexual reproduction typically focus on the benefits of the inherent ability of sexual reproduction to recombine and shuffle genetic information (*13–20*). Generally, these hypotheses have relied on fusing observational zoological studies with our understanding of genetics to build their foundations; a very understandable approach given the inherent methodological constraints on the field of evolutionary biology

Whereas hypotheses in applied physics or chemistry can be directly tested in experiments with controlled variables in real-time, evolutionary biology deals with “evolutionary time” where the timescales necessary for experimentation far exceed the human lifespan, or human civilization itself for that matter. Such controlled experiments are largely unfeasible for the field of evolutionary biology. As such, even Darwin was forced to use the inductive reasoning approach of accumulating large reservoirs of circumstantial evidence in combination with critical thinking to propound his ideas (*21*). The field of evolutionary biology has largely inherited this inductive reasoning methodology.

With the dearth of ability to conduct concrete controlled experiments to prove evolutionary principles by a deductive approach, we believe that turning back to a mathematical approach, as done by such past luminaries as JBS Haldane (*22*) or Kimura and Maruyama (*23*) can be valuable for addressing the problem of sexual reproduction. We believe that reproductive biology generally follows consistent principles that are amenable to mathematical modeling; as such, a mathematical framework can be used to provide further support for existing ideas, as well as illuminate novel ideas that could be overlooked if not for a deductive mathematical approach. With a mathematical methodology in mind, we introduce several mathematical constructs to analyze population changes via sexual reproduction or asexual reproduction. We first introduce a diversity measure *DV M* defined as the number of distinct individuals/genotypes in a population divided by the total number of individuals/genotypes in the population. We derive dynamic equations of *DV M* for both asexual reproduction and sexual reproduction. We show that *DV M* increases exponentially in sexual reproduction rather than linearly as in asexual reproduction. This exponential increase in sexual reproduction’s diversity *DV M* is consistent with current hypotheses that posit genetic recombinatorial shuffling from sexual reproduction allows for more rapid adaptation.

We further introduce a diffusion measure *DF M* defined as the average “distance” of individuals in a population. Here, the distance between two individuals is defined as the percentage difference in their genes. Consequently, *DF M* measures how homogeneous a population is. We derive dynamic equations of *DF M* for both asexual reproduction and sexual reproduction. We show that, despite its diversity, sexually reproducing species are more homogeneous (smaller *DF M*) than asexually reproducing species. In other words, sexual reproduction increases diversity (by increasing *DV M*) while decreases diffusion (by reducing *DF M*). This is less intuitive, but true.

We show that a more homogeneous population has evolutionary advantages over a less homogeneous population (smaller *DF M* is better). This is because the diffusion measure *DF M* is inversely related to optimality measure *OPM* , which measures the average distance of individuals in a population to the optimal individual/genotype who is most suitable in a given environment. More precisely, we show that *OPM* = 1 − *DF M* . Therefore, a smaller *DF M* implies a larger *OPM* . Hence, a more homogeneous population is more suitable for the environment. Since sexual reproduction leads to smaller *DF M* versus asexual reproduction, it has advantage over asexual reproduction, in addition to the advantage of having a larger *DV M* .

The last performance measure we define in this paper is the survivability measure *SV M* which is the percentage of individuals having progeny. We show that the survivability measure increases as the optimality measure increases (and that survivability measure decreases as the diffusion measure increases).

By formally defining four performance measures, *DV M* , *DF M* , *OPM* , and *SV M* , we develop a formal mathematical framework, in which advantages of sexual reproduction can be precisely addressed. All the above discussions can be verified by simulations of the dynamical equations for *DV M* , *DF M* , *OPM* , and *SV M* derived from basic biology.

Some theories/hypotheses that have been proposed to address the advantages of sexual reproduction over asexual reproduction, such as genetic recombination, Muller’s Ratchet, and the Red Queen Hypothesis, can all be addressed by the mathematical framework using the performance measures proposed in this paper.

## Diversity, Diffusion, Optimality, and Survivability Measures

In this section, we present the framework of our research. We define diversity measure, diffusion (or inhomogeneity) measure, optimality measure, and survivability measure to describe a species. We also define selective pressure to describe an environment. We show how survivability measure depends on diffusion measure, optimality measure, and selective pressure.

To do this formally and mathematically, let us first introduce the following notations.

*N* - the number of genes

*g*_*i*_, 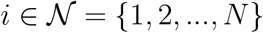 - the *i*-th gene

In asexual reproduction, a progeny inherits its genes from one progenitor.

In sexual reproduction, a progeny inherits its genes from two progenitors.

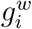 - the *i*-th gene from the maternal progenitor.

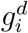 - the *i*-th gene from the paternal progenitor

A progeny will randomly pick the *i*-th gene from the maternal or paternal progenitor with equal probability (=0.5).

From one generation to the next, the probability of mutation of *g*_*i*_ is *p*.

A mutation causes *h*% change in *g*_*i*_.

Change of mutation is additive: *k* mutations cause *kh*% change in *g*_*i*_.

*Q* - the total population of a species.

*q* ∈ *Q* - an individual/genotype in the species *Q*.

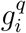 - the *i*-th gene of *q* ∈ *Q*.

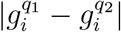 - distance between 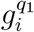 and 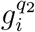, defined as the percentage difference between 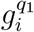 and 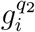.

If the distance between 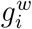 and 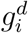 exceeds a bound *b*, that is, 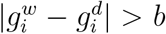, then either the mating will not take place or the resultant progeny will not survive.

### Diversity measure

Diversity measure describes how diverse a population is. We will argue that a more diverse population has an evolutionary advantage over a less diverse population, as more diversity increases the chance to adapt to the environment.

Diversity measure of a species *Q* is defined as the number of distinct individuals/genotypes in *Q* divided by the total number of individuals/genotypes in *Q*, that is,

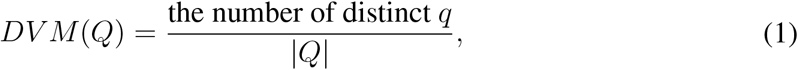

where |*Q*| is the number of elements (cardinality) of *Q*. Clearly, larger *DV M* (*Q*) means that the species is more diverse.

### Diffusion measure

Distance between *q*_1_, *q*_2_ ∈ *Q* is defined as

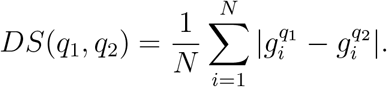

Diffusion measure of a species *Q* is defined as the average or expected value of *DS*(*q*_1_, *q*_2_):

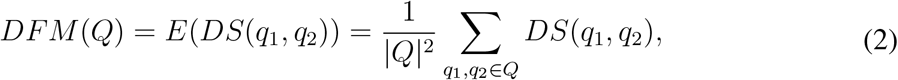

where *E*(.) denotes the expectation (mean). Clearly, larger *DF M* (*Q*) means that the species is less similar.

### Optimality measure

In a given environment, the optimal individual is denoted by *o* ∈ *Q*, whose gene is denoted by 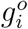, 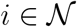 . The goal of evolution is for the population to evolve around *o*, that is, the distance of *Q* to *o* reduces as time passes.

The distance of *Q* from *o* is defined as the average or expected distance of individuals *q* ∈ *Q* to *o*. One minus this distance is called the optimality measure *OPM* . Hence, optimality measure of a species *Q* is one minus the expected value of *DS*(*q, o*):

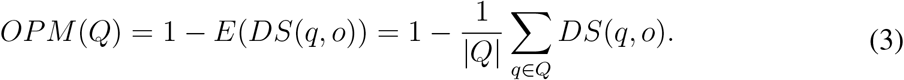

The reason to use 1 − *E*(*DS*(*q, o*)) rather than *E*(*DS*(*q, o*)) as optimality measure is that, intuitively, we would like to see that the larger *OPM* is, the closer *Q* is to *o*.

As to be shown in Section V, optimality measure is related to diffusion measure as follows.

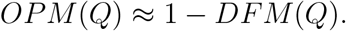

### Selective pressure

For a given environment, its selective pressure is measured as a threshold *SP* ∈ [0, 1] such that, if *DS*(*q, o*) > 1 − *SP* , then individual *q* will not have progeny. The larger *SP* is, the strong selective pressure is.

### Survivability measure

In a given environment, survivability measure of a species *Q* is defined as the percentage of individuals having progeny, that is,

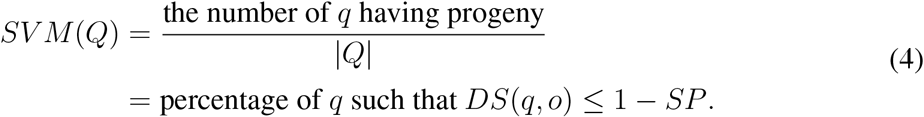

Survivability measure depends on selective pressure, optimality measure, and diffusion measure as shown in the following theorem, whose proof is in Appendix.

#### Theorem 1

(1) *As the selective pressure SP increases, the survivability measure SV M* (*Q*) *decreases.*

(2) *As the optimality measure OPM* (*Q*) *increases, the survivability measure SV M* (*Q*) *increases.*

(3) *As the diffusion measure DFM* (*Q*) *increases, the survivability measure SV M* (*Q*) *decreases.*

In the next two sections, we derive the diversity and diffusion measures for both asexual reproduction and sexual reproduction. Without loss of generality, we assume that all individuals in *Q* are of the same generation. Denote the *m*-th generation of the species as *Q*_*m*_. Since *DV M* (*Q*), *DF M* (*Q*), *OPM* (*Q*), and *SV M* (*Q*) are all defined as a percentage, we further assume that, without loss of generality, the number |*Q*_*m*_| of individuals in *Q*_*m*_ is unchanged from one generation to the next.

### Asexual Reproduction

For asexual reproduction, each individual in the m-th generation *q*′ ∈ *Q*_*m*_ is produced by an individual in the m-1-th generation *q* ∈ *Q*_*m*−1_, denoted as *q* → *q′*.

#### Diversity measure

For asexual reproduction, each mutation produces an individual who is different from its parent. The probability that this individual is distinct in *Q* is proportional to 1 − *DV M* (*Q*_*m*−1_). The expected number of mutations in one generation is given by |*Q*| × *N* × *p*. Hence,

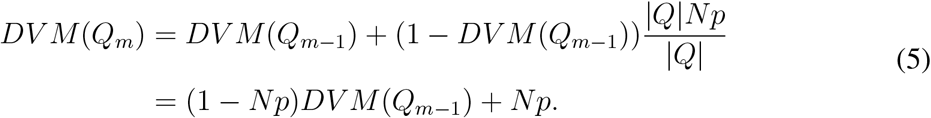

It can be shown (Section V) that *DV M* (*Q*_*m*_) → 100% as *m* → ∞. Assume that *DV M* (*Q*_*m*_) is small initially, then 1 − *DV M* (*Q*_*m* − 1_) ≈ 1 and the above equation can be approximated as

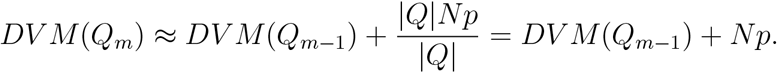

In other words, *DV M* (*Q*_*m*_) increases linearly.

#### Diffusion measure

To calculate *DS*(*q*′_1_, *q*′_2_), where *q*_1_ → *q*′_1_ and *q*_2_ → *q*′_2_, let us consider their *i*-th genes, 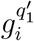 and 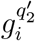. Since mutations are independent with probability (w.p.) *p*, we have the following

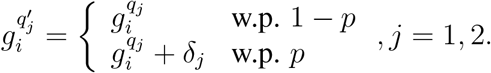

where *δ* is used to denote a mutation. Hence, the possible pairs 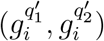 and probabilities of their occurrences are as follows.

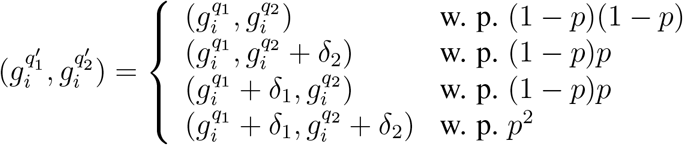

A mutation causes *h*% change. The probability that this mutation increases 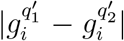 is proportional to 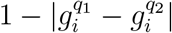 . Thus,

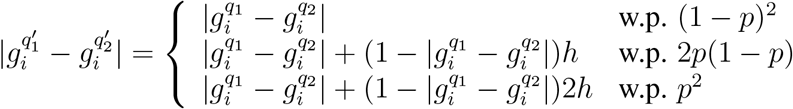

Hence^1^,

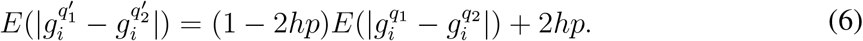

Therefore,

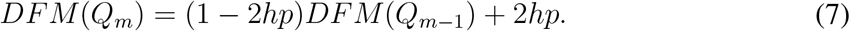

It can be shown (Section V) that *DF M* (*Q*_*m*_) → 100% as *m* → ∞. In other words, in asexual reproduction, individuals in a species will become less and less similar as time passes.

### Sexual Reproduction

In sexual reproduction, each individual in the m-th generation *q*′ ∈ *Q*_*m*_ is produced by two individuals in the m-1-th generation *q*_*w*_, *q*_*d*_ ∈ *Q*_*m*−1_, denoted as *q*_*w*_, *q*_*d*_ → *q′*.

#### Diversity measure

While in asexual reproduction, one new mutation will increase the number of distinct new individuals by 1, in sexual reproduction, one new mutation has the potential to double the number of distinct new individuals due to genetic recombination. We assume that *c* percentage of genetic recombination will be accomplished. Since the expected number of mutations in one generation is |*Q*| × *N* × *p*, the expected number of new individuals due to genetic recombination is |*Q*|*cNp* × *DV M* (*Q*_*m*−1_)|*Q*|. The probability that the new individuals are distinct in *Q*_*m*_ is proportional to 1 − *DV M* (*Q*_*m*−1_). Hence,

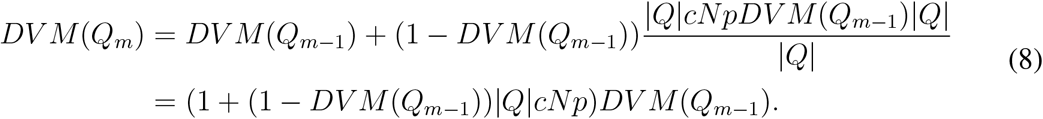

It can be shown (Section V) that *DV M* (*Q*_*m*_) → 100% as *m* → ∞. Assume that *DV M* (*Q*_*m*_) is small initially, then 1 − *DV M* (*Q*_*m*−1_) ≈ 1 and the above equation can be approximated as

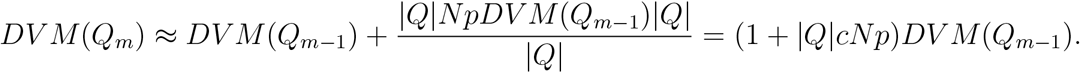

In other words, *DV M* (*Q*_*m*_) increases exponentially with respect to *m*. It is well-known that exponential increase is much faster than linear increase.

#### Diffusion measure

To calculate *DS*(*q*′_1_, *q*′_2_), where *q*_1,*f*_, *q*_1,*d*_ → *q*′_1_ and *q*_2,*f*_ , *q*_2,*d*_ → *q*′_2_ , let us consider their *i*-th gene, 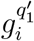 and 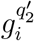. Since mutations are independent with probability *p*, we have

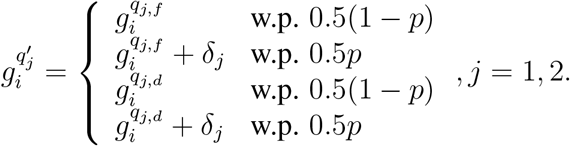

A mutation causes *h*% change. The probability that this mutation increases 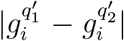 is proportional to 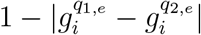 , where *e* ∈ {*f, d*}. Thus,

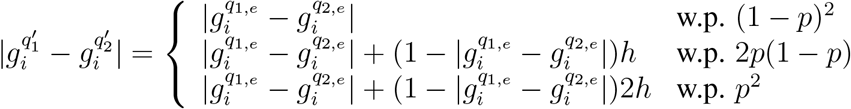

Since, 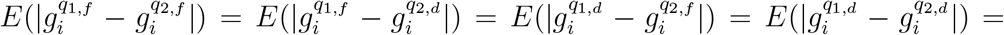 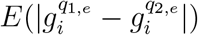, let us denote them by 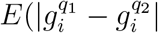. Then,

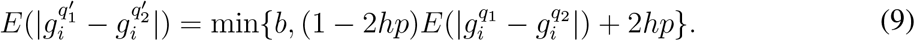

The reason for min is that, if 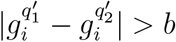, then the progeny will not survive. Therefore,

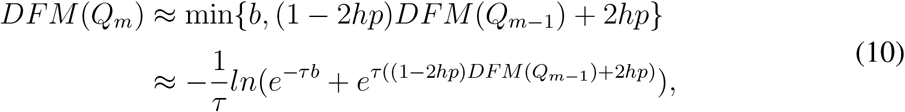

where *τ* > 0 is the parameter that determines the softness of the approximation of min.

It can be shown (Section V) that *DV M* (*Q*_*m*_) → *b* as *m* → ∞. This is very different than asexual reproduction, where *DF M* (*Q*_*m*_) → 100% as *m* → ∞. In other words, while asexual reproduction can make individuals in a species completely dissimilar, to the point that a taxonomist could be forced to concede that the population of individuals are in fact multiple “species,” whereas, sexual reproduction maintains a species as a related and coherent group.

### Simulation Results

There are two tasks for simulations. The first task is to verify

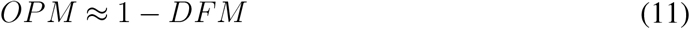

The second is to compare sexual reproduction with asexual reproduction by means of the diversity, diffusion, and optimality measures.

We use MATLAB, a popular engineering software to perform simulations. For the first task, we calculate *DF M* and *OPM* using Equations (2) and (3).

Since *DS*(*q*_1_, *q*_2_) and *DS*(*q, o*) are percentages, we represent *q* ∈ *Q* by a number between 0 and 1. We generate *K* random numbers, denoted, with a slight abuse of notation, as *Q* = {*q*_1_, *q*_2_, …, *q*_*K*_}, and then calculate

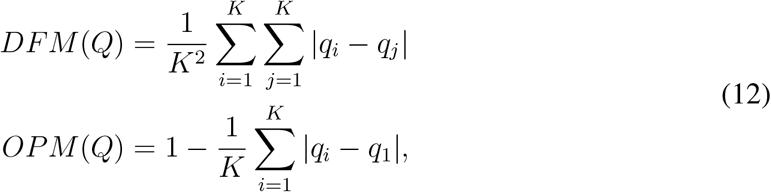

where, without loss of generality, we let *o* = *q*_1_.

We run the above simulations *R* times and take the average. For different *K* and *R*, the results are shown in Table 1.

**Table 1:** Simulation results to verify *OPM* ≈ 1 − *DF M* or *OPM + DFM* ≈ 1.

| K | R | DFM | OPM | SUM | ERROR |
| --- | --- | --- | --- | --- | --- |
| 3000 | 200 | 0.3331 | 0.6679 | 1.0010 | 0.1% |
| 1000 | 100 | 0.3337 | 0.6610 | 0.9947 | -0.5% |
| 1000 | 300 | 0.3331 | 0.6716 | 1.0047 | 0.5% |
| 5000 | 100 | 0.3336 | 0.6590 | 0.9926 | -0.7% |
| 5000 | 300 | 0.3332 | 0.6683 | 1.0015 | 0.2% |

For the second task, we simulate the derived dynamic equations, which are summarized below, to compare sexual reproduction with asexual reproduction. Since it is a comparison, only the relative values (rather than the absolute values) of the diversity, diffusion, and optimality measures are of importance.

#### Diversity measure

Diversity measure for asexual reproduction is given in Equation (5). Diversity measure for sexual reproduction is given in Equation (8).

#### Diffusion measure

Diffusion measure for asexual reproduction is given in Equation (7). Diffusion measure for sexual reproduction is given in Equation (10).

#### Optimality measure

Diffusion measures for both asexual and sexual reproductions are given by Equation (11).

We conducted several simulations with different parameter values. The results are all similar. Typical parameters used in the simulations are as follows. We do not claim that these parameters are biologically representative. Note that since we are only interested in the relative values of the diversity, diffusion, and optimality measures, absolute values of parameters are not critical.

*p* = 0.001, *h* = 25%, *N* = 10, |*Q*| = 200, *b* = 30%, *c* = 10%, *τ* = 10.

The initial conditions are *DV M* (*Q*_0_) = 10% and *DF M* (*Q*_0_)) = 10%. Results of simulations are shown in Figures 1 and 2.

**Figure 1:**
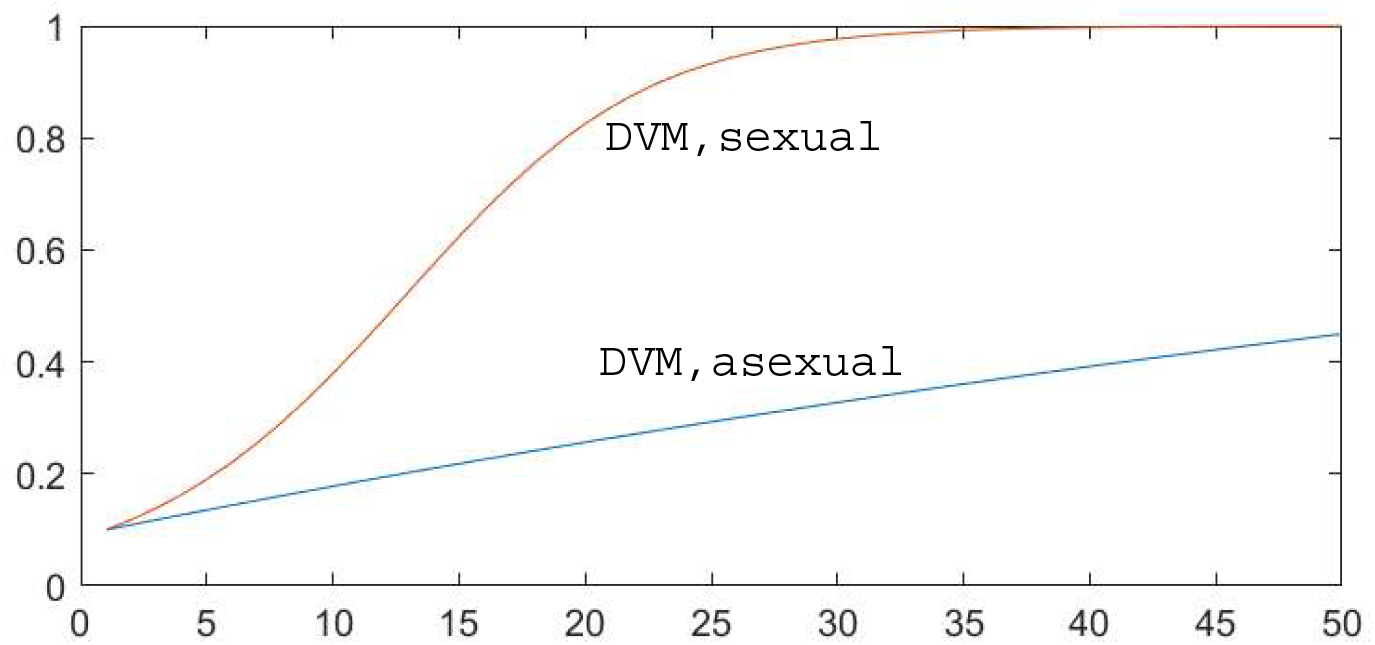
Diversity measure *DV M* of sexual vs asexual reproduction. Horizontal axis shows the number of generations.

**Figure 2:**
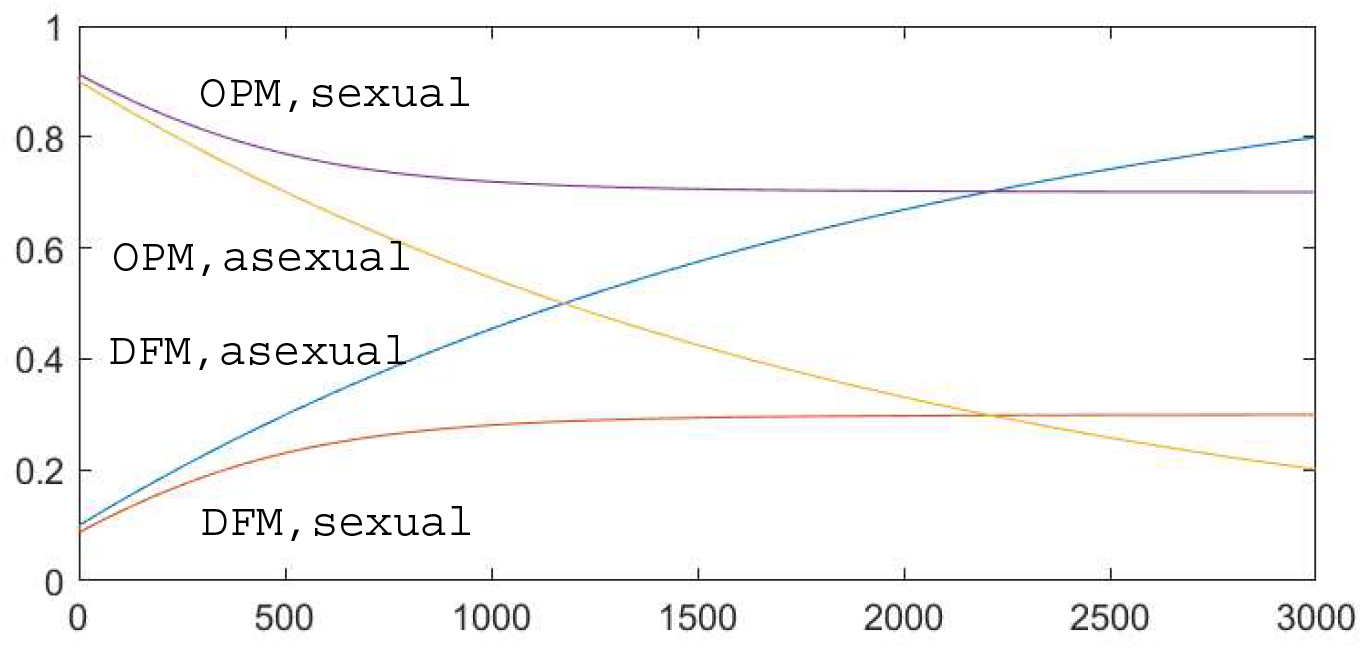
Diffusion measure *DF M* and optimality measure *OPM* of sexual vs asexual reproduction. Horizontal axis shows the number of generations.

From Figure 1, it is clear that the diversity measure of sexual reproduction is larger than and increases faster than that of asexual reproduction. Larger diversity measure allows a species to evolve faster. Hence, in terms of diversity measure, sexual reproduction has an advantage over asexual reproduction.

On the other hand, from Figure 2, diffusion measure of sexual reproduction is bounded and smaller than that of asexual reproduction. By Theorem 1, smaller diversity measure leads to larger survivability measure. Hence, in terms of diffusion measure, sexual reproduction also has an advantage over asexual reproduction.

Comparing the horizontal axis of Figure 1 with the horizontal axis of Figure 2, we note that the advantage of sexual reproduction over asexual reproduction in terms of diffusion measure takes a longer time to show than the advantage in terms of diversity measure. This probably explains why the effects of diversity measure have been so readily latched onto and investigated by biologists, whereas the potential evolutionary consequences of diffusion measure’s effects have been overlooked and not been investigated in any comprehensive manner.

### Diversity vs Diffusion

By the definitions, *DV M* is the number of distinct individuals in a population *Q* divided by the total number of individuals in *Q*, while *DF M* is the average distance between any two individuals in *Q*. Hence, *DV M* describes how diverse a population is, while *DF M* describes how similar or homogenous a population is. At first glance, one may think that *DV M* and *DF M* depend on each other. However, they are actually independent of each other. This is illustrated in Figure 3, where two populations have the same *DV M* , but the population on the left has a much smaller *DF M* that the one on the right. Hence, it is entirely possible to keep *DV M* large while keeping *DF M* small. We argue that, for a population to best adapt to an environment, it is best for the population *Q* to have a large *DV M* and a small *DF M* (see Theorem 1).

**Figure 3:**
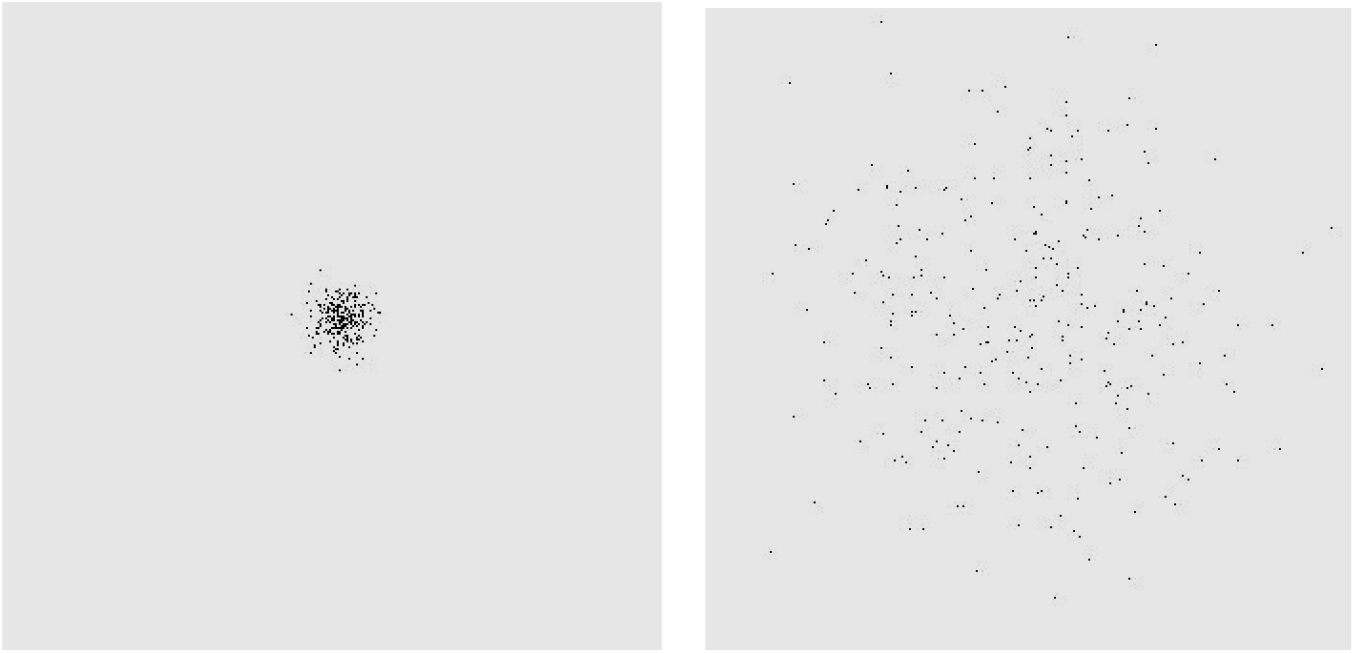
Diversity measure *DV M* and diffusion measure *DF M* are independent of each other. The above two populations have the same *DV M* , but the population on the left has a much smaller *DF M* than the one on the right.

Sexual reproduction allows genetic recombination to produce recombinant types via mechanisms such as chromosome shuffling, and this process occurs with each successive progeny. Subsequently, through this probabilistic process, every progeny is forced to be distinct from both its progenitors and sibling progeny (assuming the progenitors do not have exactly identical genomes, and excluding atypical phenomena such as twins, etc.). In other words, sexual reproduction results in a high *DV M* . This point has been well established. Our added contribution is the insight that such a chromosomal reshuffling phenomenon could only occur if both progenitors are not radically dissimilar, such as a paternal progenitor with 46 chromosomes contributing 23 chromosomes and a maternal progenitor with 44 chromosomes contributing 22 chromosomes. This is an example of a high *DF M* value precluding the generation of progeny. Asexual reproduction has no such constraints. A progenitor of 46 chromosomes could produce 10 progenies: 9 with identical 46 chromosomes, and 1 with 47 chromosomes as a mutational accident. Such a population would have low *DV M* (90% of the population are 100% identical) with high *DF M* (10% of the population is radically different in chromosome number).

Our mathematical models and simulation results are consistent with the evidences that sexual reproduction promotes genetic similarity/homogeneity, while asexual reproduction leads to genetic diffusion. As seen from our simulations, asexual reproduction results in an increase of *DF M* over time, while sexual reproduction results in bounded *DF M* , implying the formation of a tight cluster of similar individuals. The tight cluster is maintained by sexual reproduction that prevents the diffusion seen in asexual reproduction. Theorem 1 states that a smaller *DF M* means a large *OPM* and hence a large *SV M* .

To illustrate this benefit of genetic similarity and maintenance of adaptational advantages in a biological context, we take the example of a developed ecosystem that is near its “climax community”. Such a system can be considered relatively stable with high biodiversity. The high biodiversity means there is more competition for the same limited resources. In order for all these species to successfully live amongst each other, selective pressures have caused each species to develop specific adaptations that allow it to occupy an exclusive niche. Deviating away from such adaptations, which means deviating away from the species’ niche, results in competition with other species that occupy other niches. These other species are highly adapted for their niches; consequently, the “deviant organism” is unlikely to survive the competition. Sexual reproduction maintains adaptational advantages and minimizes the conversion of precious resources to the production of deviant organisms that are unlikely to survive.

In summary, both *DV M* and *DF M* are distinct principles that can provide a more nuanced approach to investigating the advantages of sexual reproduction over asexual reproduction. (1) Sexual reproduction leads to a more diverse population and hence allows more rapid adaption to environments. (2) Sexual reproduction is a species stabilization mechanism that naturally maintains genetic homogeneity and species identity. (3) Asexual reproduction, which does not have this inherent species stabilization mechanism, leads to genetic inhomogeneity and no definitive species identity. (4) Sexual reproduction is beneficial because the maintenance of species identity maintains desired adaptational advantages, which is important when selective pressures are strong. In the next section, we use *DV M* and *DF M* to explain some theories/hypotheses that have been proposed to address the advantages of sexual reproduction over asexual reproduction.

### Advantages of Sexual Reproduction

Various theories/hypotheses have been proposed to address the advantages of sexual reproduction over asexual reproduction. The main theories/hypotheses are as follows. (1) *Genetic recombination:* Sexual reproduction allows genetic recombination to produce recombinant types that can make the population better able to adapt changes in the environment. In other words, sexual reproduction allows greater exploration of genotype possibilities hence providing the raw fuel for natural selection to act upon. (2) *Muller’s Ratchet:* Continued accumulation of deleterious mutations leads to a degradation of a species’ average fitness over time, without any method of correction, the species will likely face extinction (*24–26*). Muller’s Ratchet posits that sexual reproduction allows good genes in separate loci to be recombined to restore genotypes to optimal fitness. (3) *Red Queen hypothesis:* In this hypothesis, parasites and hosts engage in a constant arms battle with hosts evolving resistances to parasites and parasites evolving ways to get past those resistances (*10, 27, 28*). Sexual reproduction generates new genotypes at a much faster rate than would be possible with asexual reproduction. The faster generation of new genotypes then allows for rapid adaptation of resistances against parasites for the hosts. Likewise, sexually reproducing parasites can have rapid adaptation against host resistances. The mathematical models we presented fits well with these theories/hypotheses.

1. *Genetic recombination:* The diversity measure *DV M* describes genotype possibilities of a population. Comparing Equation (5) with Equation (8), we can see that initially (when *DV M* is small), *DV M* increases linearly in asexual reproduction, but exponentially in sexual reproduction. Simulation results of Figure 1 clearly show that *DV M* increases much faster in sexual reproduction than in asexual reproduction.
2. *Muller’s Ratchet:* Muller’s Ratchet is reflected in the optimality measure *OPM* . As shown in Figure 2, *OPM* decreases in both sexual reproduction and asexual reproduction as time goes by. This is exactly what Muller’s Ratchet anticipates. However, in sexual reproduction, *OPM* will reach a limit and no longer decreases. In other words, sexual reproduction can restore genotypes to optimal fitness.
3. *Red Queen hypothesis:* The Red Queen Hypothesis can also be explained by *DV M* . Sexually reproduced parasites and hosts have much larger *DV M* than asexually reproduced parasites and hosts as shown in Figure 1. Larger *DV M* allows parasites and hosts to better adapt in the battle with hosts evolving resistances to parasites and parasites evolving ways to get past those resistances.

## 1 Appendix Proof of Theorem 1

Let us enumerate individuals in *Q* based on their distance to *o*, that is, let

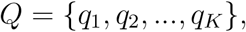

where *K* = |*Q*|, such that

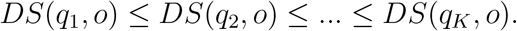

Define a distance function 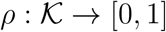, where 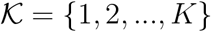, as

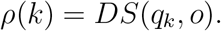

Then *ρ* is a monotonic increasing function, that is,

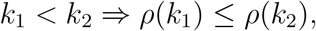

where ⇒ denotes “implies”.

From the definition of survivability, we have

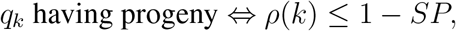

where ⇔ denotes “if and only if”.

Therefore,

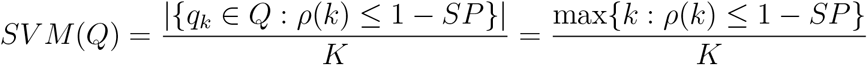

To prove Part (1) of Theorem 1, consider two selective pressures *SP*_1_, *SP*_2_ with *SP*_1_ ≤ *SP*_2_.

We have

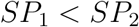

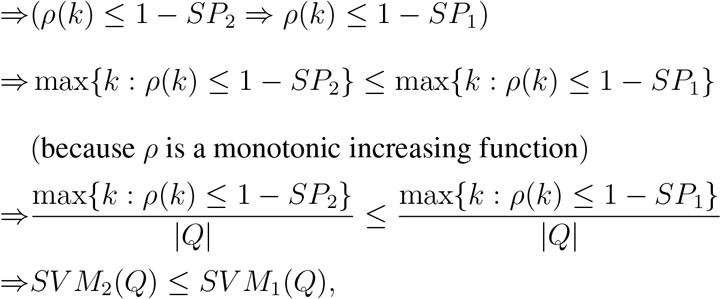

where *SV M*_*i*_(*Q*) is the survivability measure under selective pressure *SP*_*i*_, *i* = 1, 2.

To prove Part (2) of Theorem 1, consider two optimality measures *OPM* (*Q*_1_), *OPM* (*Q*_2_) with *OPM* (*Q*_1_) ≤ *OPM* (*Q*_2_). By the definition of *OPM* (*Q*_*i*_), *i* = 1, 2,

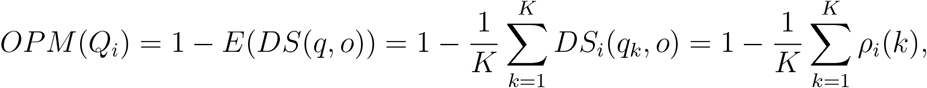

where *ρ*_*i*_(*k*) = *DS*_*i*_(*q*_*k*_, *o*), for 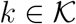, corresponds to *OPM* (*Q*_*i*_). Hence, for any *SP*

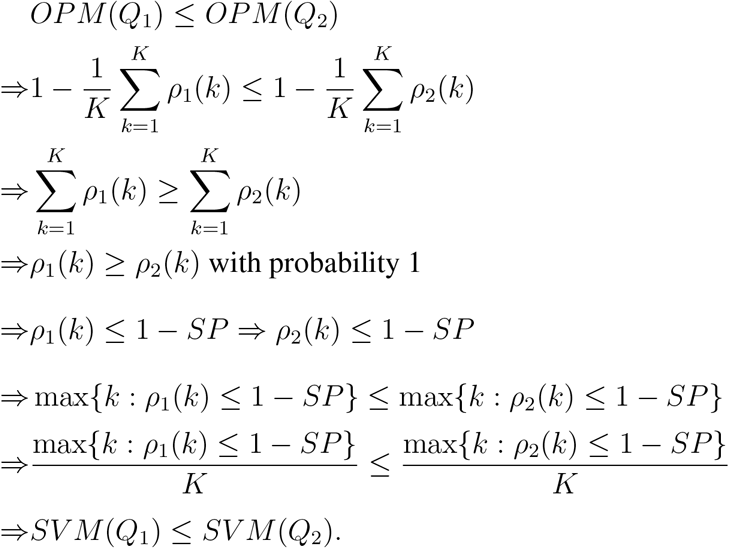

Part (3) of Theorem 1 is the direct result of Part (2) of Theorem 1 and the relation *OPM* (*Q*) ≈ 1 − *DF M* (*Q*).

Derivation of Equation (6)

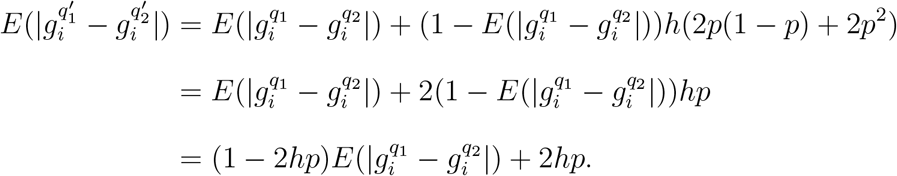

Derivation of Equation (7)

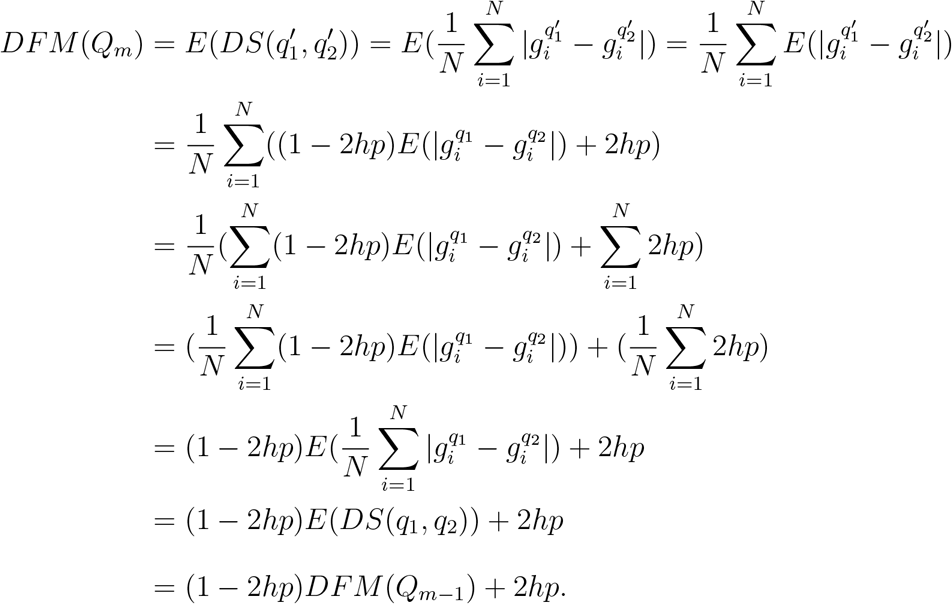

Derivation of Equation (9)

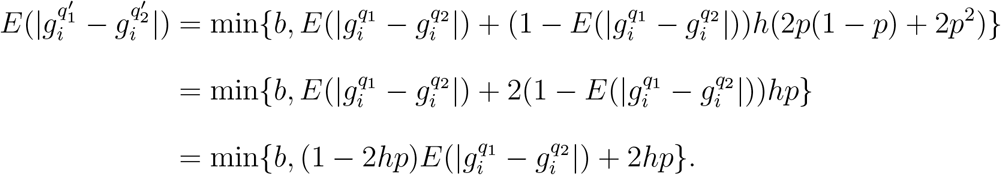

Derivation of Equation (10)

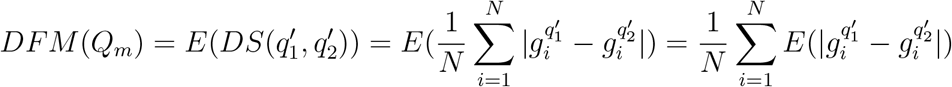

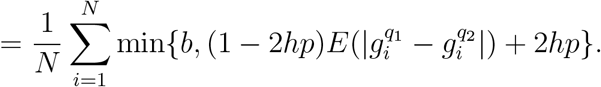

Mathematically,

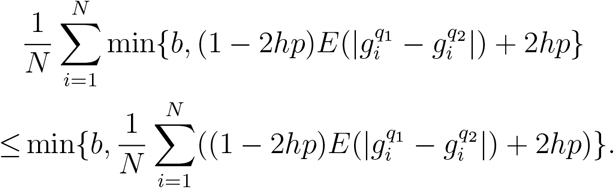

However, we expect that 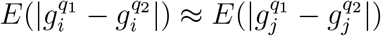, for 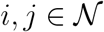 . Therefore,

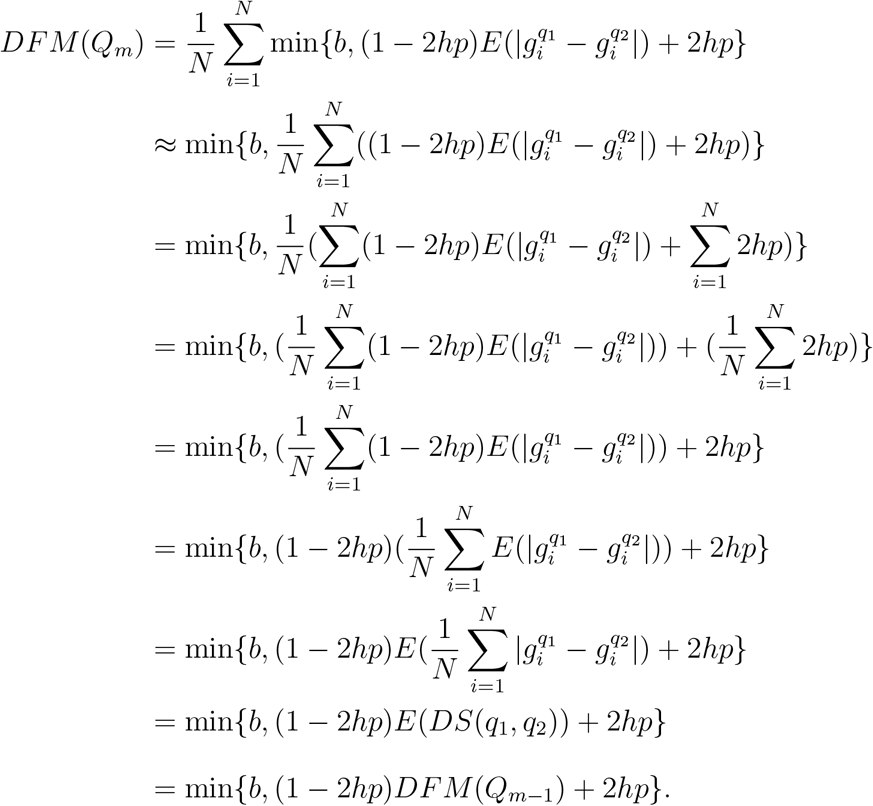

For better simulations, we approximate min by a continuous function as follows.

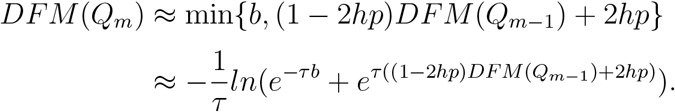

## Footnotes

1 Derivations of all equation can be found in Appendix.

